# Dissociable causal roles of dorsolateral prefrontal cortex and primary motor cortex as a function of motor skill expertise

**DOI:** 10.1101/2023.10.20.563280

**Authors:** Quynh N. Nguyen, Katherine J. Michon, Taraz G. Lee

## Abstract

Established models of motor skill learning posit that early stages of learning are dominated by an attentionally demanding, effortful mode of control supported by associative corticostriatal circuits involving the dorsolateral prefrontal cortex (DLPFC). As expertise develops, automatic and “effortless” performance coincides with a transition to a reliance on sensorimotor circuits that include primary motor cortex (M1). However, the dynamics of how control evolves during the transition from novice to expert are currently unclear. This lack of clarity is due, in part, to the fact that most motor learning studies comprise a limited number of training sessions and rely on correlative techniques such as neuroimaging. Here, we train human participants on a discrete motor sequencing task over the course of six weeks, followed by an assessment of the causal roles of DLPFC and M1 at varying levels of expertise. We use repetitive transcranial magnetic stimulation to transiently disrupt activity in these regions immediately prior to performance in separate sessions. Our results confirm the dissociable importance of DLPFC and M1 as expertise develops. DLPFC stimulation leads to larger behavioral deficits for novice skills than expert skills, while M1 stimulation leads to relatively larger deficits as expertise develops. However, our results also reveal that prefrontal disruption causes performance deficits at all levels of expertise. These findings challenge existing models and indicate an evolving rather than a strictly diminishing role for DLPFC throughout learning.

**Significance Statement:** Motor skills involve the sequential chaining of individual actions. For example, playing the piano involves learning to rapidly transition to from one finger press to another. Human neuroimaging studies have shown that primary motor cortex (M1) and dorsolateral prefrontal cortex (DLPFC) support novice motor sequencing skills, but activity in both regions declines over training. This has been interpreted as increased efficiency in M1 and yet a reduction in the involvement of DLPFC as expertise develops. We causally test this assumption by using non-invasive brain stimulation to transiently disrupt cortical activity following extended skill training. Although we confirm dissociable contributions of DLPFC and M1 as expertise develops, we show that both regions are necessary for performance regardless of skill level.

## Introduction

Most models of motor skill learning suggest that with practice, movements transition from being attentionally demanding and effortful to more automatic (Abrahamse et al., 2013; Fitts & Posner, 1967; Newell, 1991; Willingham, 1998). The neurophysiological changes that accompany motor expertise development are often studied by examining motor sequencing tasks that involve merging motor primitives together to create a smooth, multi-component action. These tasks are intended to mimic complex actions, such as a tennis serve, that require binding serial actions together. During the early stages of learning, performing motor skills is thought to require cognitive resources from an associative corticostriatal circuit that includes the dorsolateral prefrontal cortex (DLPFC; Doyon & Benali, 2005; Hikosaka et al., 2002; Verwey et al., 2019) This associative circuit is thought to be released from control as a ‘slow-learning’ sensorimotor corticostriatal circuit, including primary motor cortex (M1), dominates performance at later stages of learning (Doyon & Benali, 2005; Hikosaka et al., 2002).

However, recent work has questioned the roles of DLPFC and M1 in skilled motor performance. Functional magnetic resonance imaging (fMRI) studies suggest that M1 is responsible for executing individual discrete movements rather than containing skill-specific representations (Berlot et al., 2020; Yokoi et al., 2018; Yokoi & Diedrichsen, 2019). Further, M1 may not be necessary for the expression of an expert skill at all, as its ablation in rodents does not alter performance following extensive training (Kawai et al., 2015). Several neuroimaging and non-invasive brain stimulation studies suggest that increased activity in the DLPFC may be detrimental to the long-term development of expertise and performance following initial training (Bassett et al., 2015; Cohen & Robertson, 2011; Galea et al., 2010). A fast pace of improvement over the course of training may rely on rapid reductions in the contribution of DLPFC (Bassett et al., 2015) and using transcranial magnetic stimulation (TMS) to disrupt DLPFC activity just following an initial training session has been shown to result in improved performance in subsequent sessions (Cohen & Robertson, 2011; Galea et al., 2010).

Neural plasticity likely involves multiple, temporally overlapping processes, complicating our understanding of how the brain supports motor skill learning. For example, learning leads to a shift in neuronal recruitment whereby new neuronal populations become engaged(Dayan & Cohen, 2011; Doyon & Benali, 2005; Wymbs & Grafton, 2015). Other brain regions that were integral to performance early on show reductions in activity as measured by fMRI because they are no longer needed(Toni et al., 1998; Ungerleider, 2002; Wymbs & Grafton, 2015). However, reduced activity can also reflect increases in efficiency whereby neurons maintain the same function while receiving less presynaptic input (Picard et al., 2013). These diverging possibilities have made inferences from studies using fMRI difficult, as there is evidence for both increased and decreased activity levels in various regions of interest (Dayan & Cohen, 2011; Doyon & Benali, 2005). For example, M1 and DLPFC often exhibit similar decreases in activity over the course of training (Dayan & Cohen, 2011; Steele & Penhune, 2010). However, this pattern is often interpreted in these studies as reflecting increased efficiency in M1 and a reduced role in supporting performance for DLPFC.

TMS can provide strong causal evidence for the role of brain regions in cognitive processes of interest by measuring the effect of their disruption on behavior (Pitcher et al., 2021). Most studies examining motor skill learning using TMS have examined short timescales, such as single-session training, which cannot provide insight into the long-term dynamics of neurophysiological changes necessary for skilled motor performance.

To address the shortcomings of previous studies, the present study used TMS to investigate the causal role of DLPFC and M1 in the expression of learned motor expertise in humans by studying motor sequencing skills trained over a six-week period. Our longitudinal within-participant design allowed us to assess the impact of cortical disruption on three different levels of skill in a single session. If M1 is increasingly involved in supporting skilled performance as expertise develops, we would expect disrupting M1 activity would have the most detrimental impact on highly trained skills and a relatively minimal impact on novice skills. If, however, M1 is primarily important for producing each discrete movement in isolation, we would expect a static cost to M1 disruption across all skill levels. If DLPFC is only important during the initial acquisition of motor sequencing skills and is not needed once performance nears ceiling, we would expect large deficits in the performance of novice skills following its disruption with essentially no impact on the most highly trained skills. DLPFC disruption may even lead to *improved* performance.

## Methods

### Participants

Twenty-one right-handed participants (six females; mean age: 21.7 years) were recruited locally from the University of Michigan and surrounding community. All participants gave informed consent to participate and were compensated for their time. All participants additionally underwent a TMS Adult Safety Screening to assess the potential risk of adverse reactions to TMS. All experimental protocols were approved by the Institutional Review Boards of the University of Michigan Medical School (IRBMED).

### Discrete Sequence Production task

Participants performed the DSP task using their left (non-dominant) hand (**Figure 1**). Each trial began with a color cue (1.5 s) signifying the identity of the sequence to be performed. Then a row of four gray rectangles corresponding to the four fingers of the left hand were presented at the center of the display (1s during training; variable duration 3-8s during scanning). After this delay period, one rectangle would turn white, signaling the first element of the cued sequence. As soon as participants pressed the corresponding button with the correct finger, another box would turn white according to the next element in the motor sequence and so on, until eight elements of the sequence were pressed, or until the time deadline of 3 s (8 s for training sessions). Incorrect key press at any element within a sequence would cause the corresponding rectangle to turn red and would also conclude the trial. Participants were instructed to complete each trial as quickly as possible while still being correct. For each participant, we randomly selected six different sequences from a common pool of 11 distinct sequences that were free of key repetitions and trills (e.g., 1-2-1). We randomly paired each of the selected sequences with a distinct color cue (blue, red, green, magenta, orange, purple) for each participant. This procedure ensures that the effects observed reflect differences in the level of expertise attained and do not depend on particular finger transitions or the particular sequence identity cues used.

**Figure 1.**
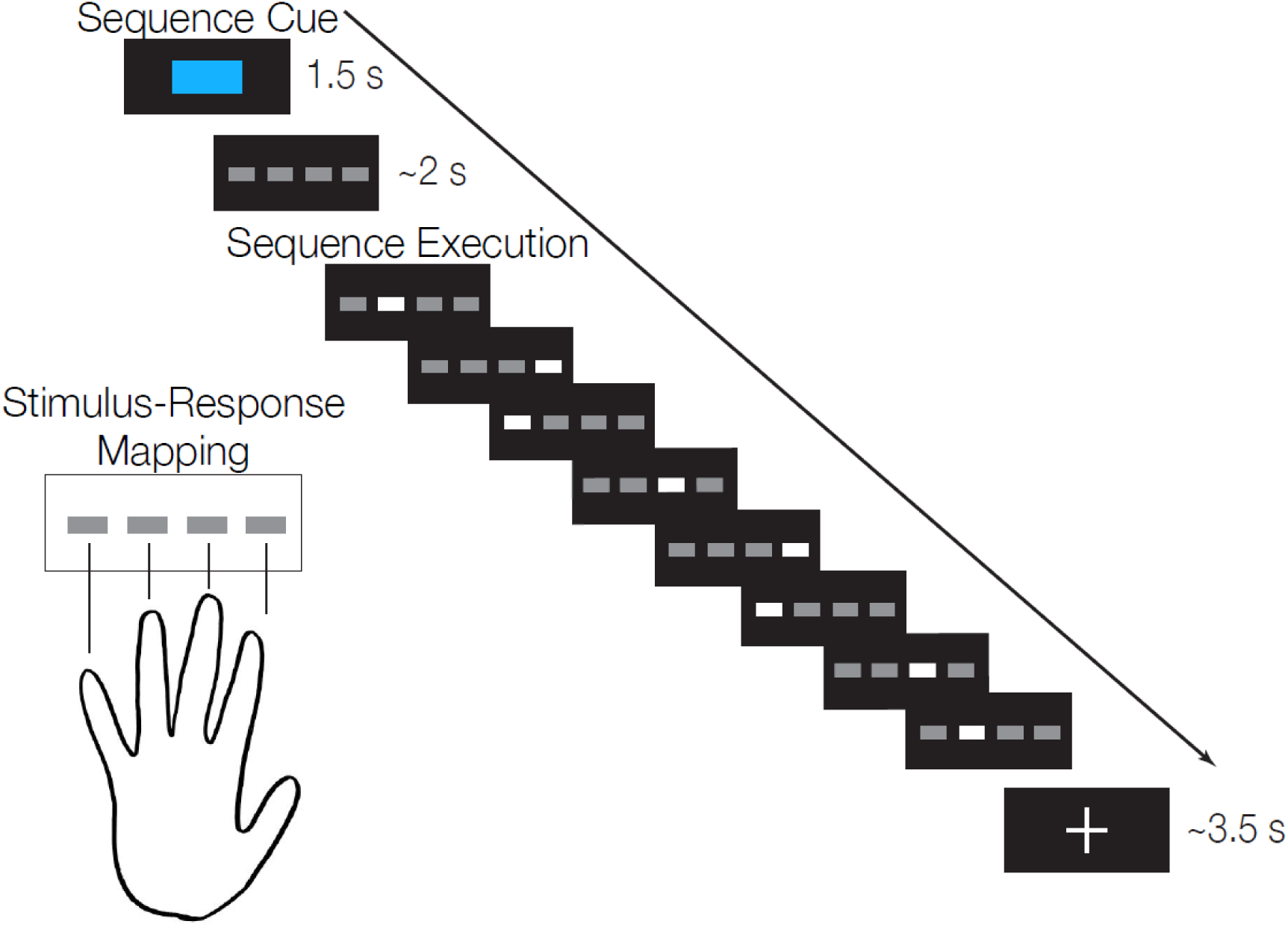
Discrete sequence production (DSP) task. On each trial, the colored cue signaled the identity of the upcoming 8-item motor sequence. Participants pressed the key corresponding to the illuminated position on the screen. As soon as a key was pressed, the next element of the sequence was immediately illuminated. The timing denoted here pertains to the baseline session and the test sessions (see Methods).

### Design overview

Following an initial screening session to determine individualized TMS parameters and assess stimulation tolerability (see TMS protocol below), each participant completed an initial fMRI scan, during which they were introduced to the discrete sequence production (DSP) task described above. Then, each participant completed 24 training sessions over a six-week period (∼4 days/week) where they practiced six different motor sequences. These sequences involved different eight-element key presses and had different practice frequencies: two sequences were practiced at 3 trials/session (minimal training; 72 trials total), another two at 15 trials/session (intermediate training; 360 trials total), and the last two at 45 trials/session (extensive training; 1080 trials total). Each training session comprised 3 blocks of 42 trials each. This training regime was designed such that each participant would obtain novice, intermediate, and expert motor skills that could be assessed concurrently in a single session following training. The first 7 participants completed training sessions on an in-lab computer. Due to the COVID-19 pandemic, training for the remaining 14 participants was completed on their personal computers at home to minimize in-person contact. At the end of the training period, each participant simultaneously possessed expertise in some sequences and was a relative novice in other sequences. In the initial fMRI session, participants were not exposed to the sequences that would be minimally trained and were asked to perform the four extensive/moderate sequences for a total of 48 trials each. Participants then returned to undergo three separate sessions where inhibitory TMS (see below) was applied to either DLPFC, M1 or a control site prior to DSP performance (8 blocks of 30 trials; 40 trials of each sequence total; ∼5 minutes per block) in the fMRI scanner to assess the impact of cortical disruption on performance (see **Figure 2a** for a schematic of the overall timeline of the experiment). These sessions were spaced at least 48 hours apart to prevent carry-over effects (mean of 6.9 days between sessions; standard deviation of 3.6 days), and the site of stimulation was counterbalanced across participants. Neuroimaging data from these sessions are not reported here.

**Figure 2.**
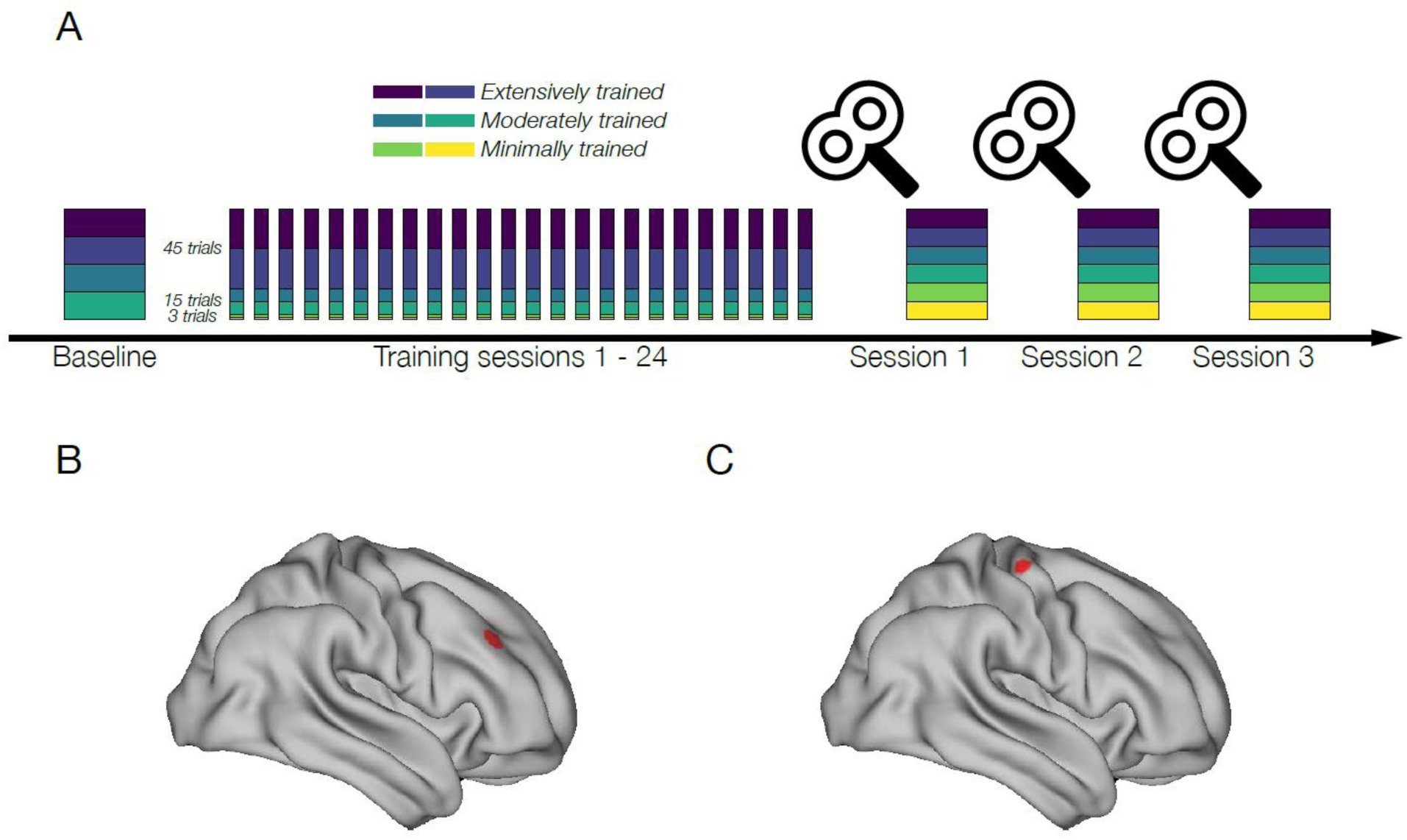
Experimental timeline. (A) Participants first attended a baseline session during which they gained initial exposure to the DSP task and some experience with the sequences that would be extensively and moderately trained. Structural MRI scans were also obtained during the baseline session. Participants then practiced the DSP task for 6 weeks (4 sessions/week) with varying training frequencies for each sequence (2 Extensive: 45 trials/session; 2 Moderate: 15 trials/session; 2 Minimal: 15 trials/session). At the end of training participants simultaneously had expertise on certain sequences but were relative novices on others. Participants then returned to 3 follow-up sessions where they received inhibitory TMS over one of three cortical targets (DLPFC, M1, or a control site) just prior to an assessment of performance on all the trained sequences. The order of stimulation was counterbalanced across participants. (B) Right dorsolateral prefrontal cortex (DLPFC) target projected onto a standard surface mesh. C) Right primary motor cortex (M1) target projected onto a standard surface mesh. Right M1 was identified anatomically for each individual. The cortical control site for stimulation (not shown) was medial post-central gyrus (putative hip/leg area of somatosensory cortex) and was likewise identified anatomically for each individual.

### TMS target definition

The right DLPFC stimulation site was defined using coordinates from a meta-analysis of executive action control (Cieslik et al., 2013). Specifically, we used a region along the right posterior inferior frontal sulcus (MNI coordinate: 37, 33, 32) that is activated by tasks involving attention to action and action execution (**Figure 2b**). Additionally, this area shows strong task-based and resting state functional connectivity with dorsal premotor cortex and pre-supplementary motor area (Cieslik et al., 2013). We created TMS targets by first normalizing each participant’s structural MRI scan to the MNI152 2009 template (Fonov et al., 2011). The resulting transformation matrix was used to reverse normalize a 5mm radius spherical region of interest (ROI) into each participant’s native space. The M1 stimulation site was defined as the hand knob of right M1 using each participant’s anatomical MRI scan (**Figure 2c**). Specifically, this site was localized anatomically for each participant based on previously reported anatomical landmarks (Yousry et al., 1997). We confirmed that a single suprathreshold TMS pulse at this location elicited a twitch in the left hand of each participant to ensure that this location was the hand area of M1 in all participants. To control for nonspecific effects of brain stimulation (cutaneous sensation, auditory stimulation, induction of current in the cortex), we selected the midline of the bilateral postcentral gyrus in each participant as a control site. This area putatively corresponds to the leg/trunk area of somatosensory cortex and was not activated by the DSP task in the baseline fMRI session. A frameless neuronavigation system (Brainsight 2, Rogue Research, Montreal CA) aligned the previously acquired structural scan to each participant’s skull in real-time to allow for precise targeting.

### TMS protocol

Repetitive TMS (rTMS) was delivered through a MagPro X100 magnetic stimulator and a 70 mm figure-8 coil (MC-B70, MagVenture Inc.). In the initial screening session, motor evoked potentials (MEP) elicited using biphasic posterior-anterior stimulation and coil oriented 45 degrees to coronal plane were recorded from left first dorsal interosseous (FDI) using surface electromyography (2-channel built in EMG device, Rogue Research, Montreal CA). Active motor threshold (AMT) was determined as the percentage of stimulator output that elicits an MEP of ≥ 200 μV peak-to-peak on five out of ten trials while the FDI muscle is contracted at 20% of maximum. At the end of this screening session, participants were given several pulses at 80% AMT over regions of the scalp roughly corresponding to DLPFC, M1, and the control site to ensure that the stimulation intensity would be tolerable in future sessions. No participants reported discomfort nor were any participants screened out in this process. In the experimental sessions following training, we delivered continuous theta-burst stimulation (cTBS) over a single region (DLPFC, M1, or the control site) with standard parameters and within consensus safety recommendations (Oberman et al., 2011; Rossi et al., 2020): 3 pulses of stimulation at 50 Hz at 80% AMT, repeated every 200 ms, for a total of 600 pulses in 40 seconds. The cTBS protocol has been found to have an inhibitory effect on the site of stimulation for approximately 60 minutes following stimulation (Huang et al., 2005), allowing an assessment of DSP task performance during the period of disruption. The mean time between the end of stimulation and the start of the first block of DSP performance was 6.83 min (SD = 1.42 min).

### Statistical analyses

For each trial, we computed the initiation time (IT), interkey intervals (IKIs), movement time (MT), and trial-level accuracy. IT is defined as the time from when the first rectangle turns white (denoting visual presentation of the first sequence element) to the time of the actual correct key press. IKI is defined as the time from any key press to the next immediate key press. MT is defined as the time from the first key press to the last key press (i.e., the total duration of all eight finger movements). IT, IKI, and MT were taken from accurate trials only. A trial is considered accurate only if all sequence elements are correctly executed within the time limit. Only trials with at least 1 key press were included in calculating accuracy rate as we assume trials with no response are due to attentional lapses rather than motor execution errors (136 trials total across all participants and all TMS sessions, ∼0.9% of all trials). Because of the main effect of training depth observed at baseline, the same absolute decrease in MT or accuracy would mean more for the extensively trained sequences that are executed very quickly than for the minimally trained sequences that are executed much more slowly. To properly weigh the impact of TMS disruption across different skill levels, we calculated the percentage change in performance relative to the control condition rather than the absolute change in MT or accuracy for each level of skill (Anderson et al., 2020).

We then assessed the effect of skill level, indexed by training depth, and of certain brain regions, indexed by TMS site, on each of the above metrics using a linear mixed-effect model. We modeled training depth and TMS site as fixed effects and participant identity as the grouping factor. We also included day order as a nuisance covariate to remove the confounding effect of extra practice on sessions involving TMS on behavioral performance. Note that figures 4-6 all plot the dependent variables with this effect of day regressed out. The full model was formulated as: *DV*_*ijk*_ = *μ* + *β*_*skil*_*skill*_*j*_ + *β*_*site*_*site*_*k*_ + *β*_*skil*×*site*_*skill*_*j*_*site*_*k*_ + *β*_*day*_*day* + *γ*_*i*_ + *ε*_*ijk*_, for participant *i* performing a sequence previously trained to skill level *j* immediately after receiving TMS to site *k*. *DV* can be any of the behavioral metrics defined above. We used trialwise *DV* values for all analyses, except for vertex-day accuracy due to non-constant residuals in the model fit. We remedied this heteroscedasticity by averaging the accuracy rate per sequence per subject and then taking the logit transformation of the average. We implemented all models in python using the statsmodels package (Seabold & Perktold, 2010).

**Figure 3.**
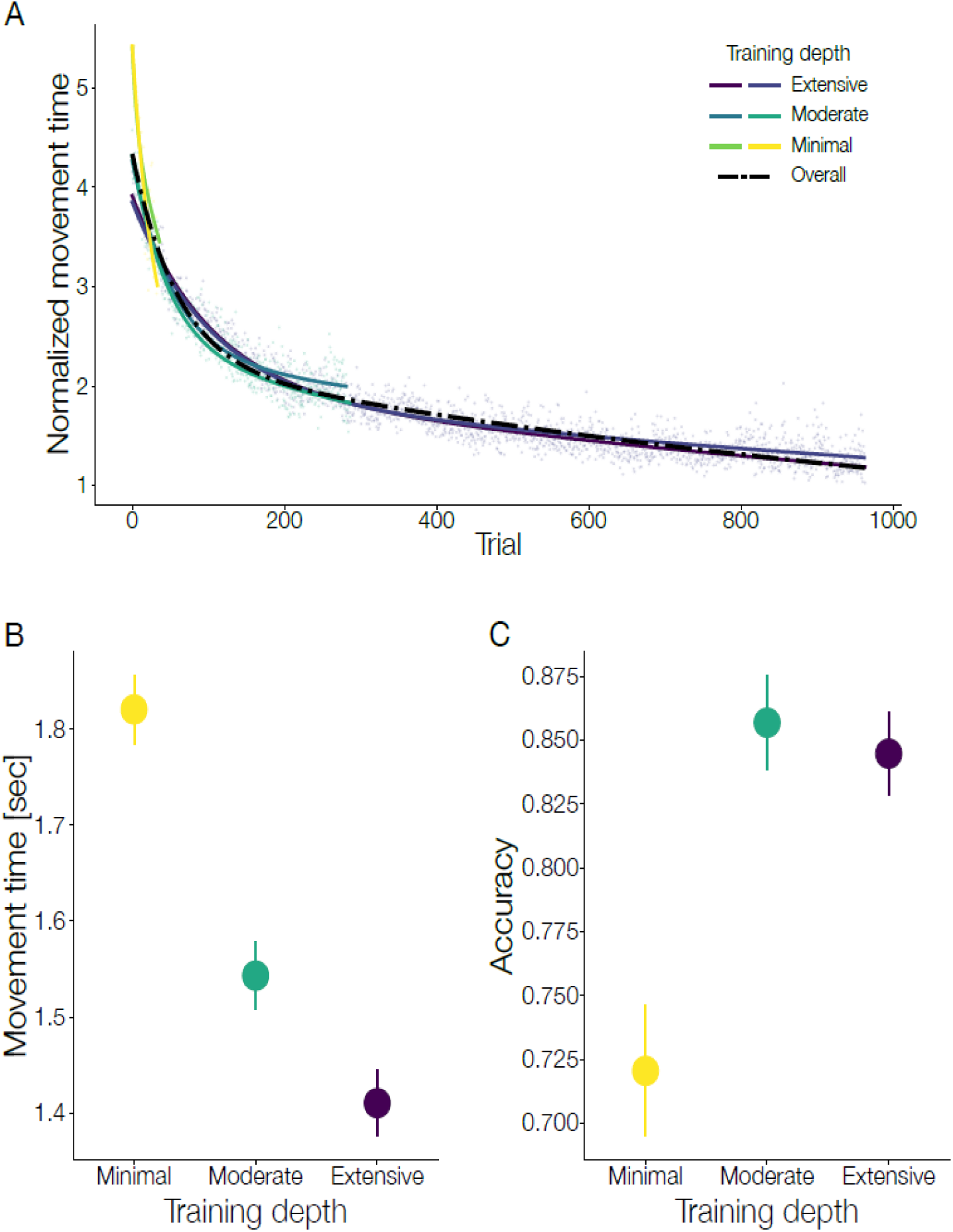
DSP performance over the course of six weeks of training. (A) Movement time (MT) decreased universally and uniformly across all sequences. Curves denote double-exponential curves fit to the group-average MT for each sequence separately (colored lines) and combined across sequences (dotted line). Note the overlapping learning trajectories. (B) After training was complete, participants exhibited the desired gradient of skill level where minimally trained sequences were executed the most slowly, moderately trained sequences somewhat more quickly, and extensively trained sequences the most quickly (data from control stimulation session). (C) The difference in execution speed is not due to a speed-accuracy tradeoff across skill level as minimally trained sequences were the least accurately performed and moderately and extensively trained sequences showed similar levels of accuracy. Note the data in B and C reflect raw movement time and accuracy for visualization rather than the standardized measures used in the analyses. Error bars represent SEM.

**Figure 4.**
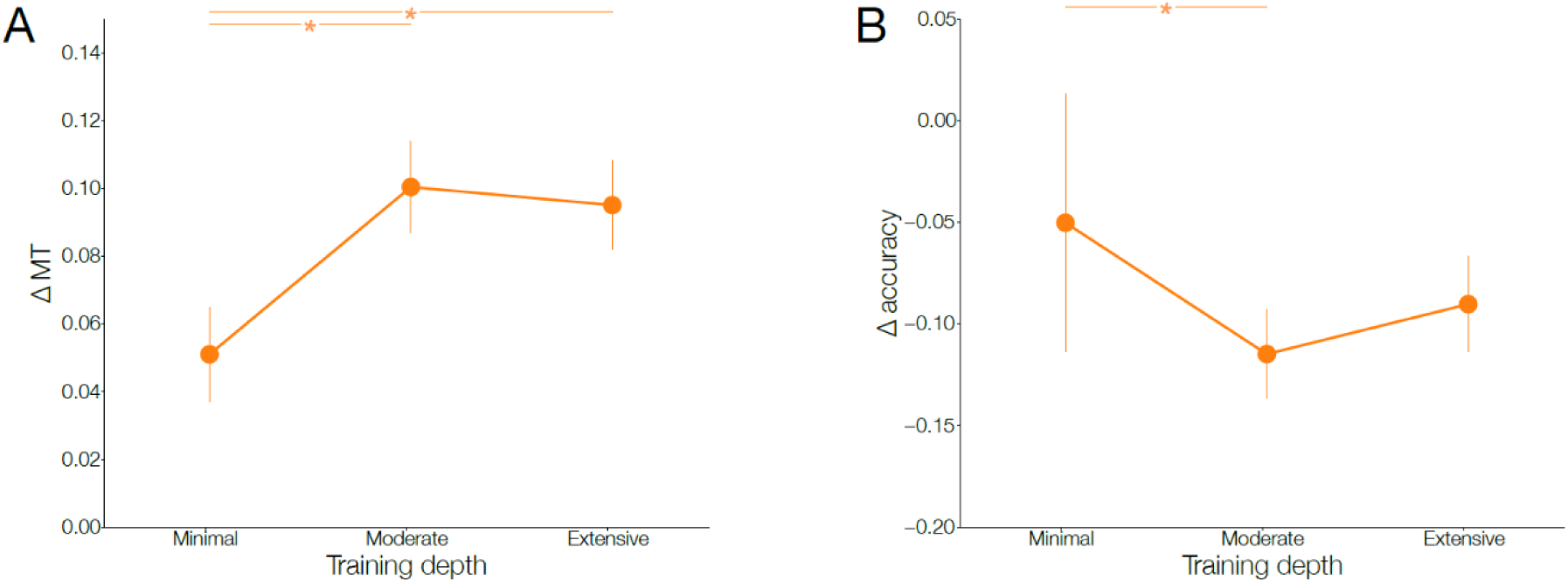
Performance changes following M1 stimulation. M1 disruption led to deficits in both (A) the speed of execution and (B) the accuracy of execution relative to performance after control stimulation. However, this performance cost was graded by skill level such that the minimally trained skills were impacted the least relative to the moderately and extensively trained skills. ΔMT = % change in movement time relative to control stimulation. Δ accuracy = % change in accuracy relative to control stimulation.

**Figure 5.**
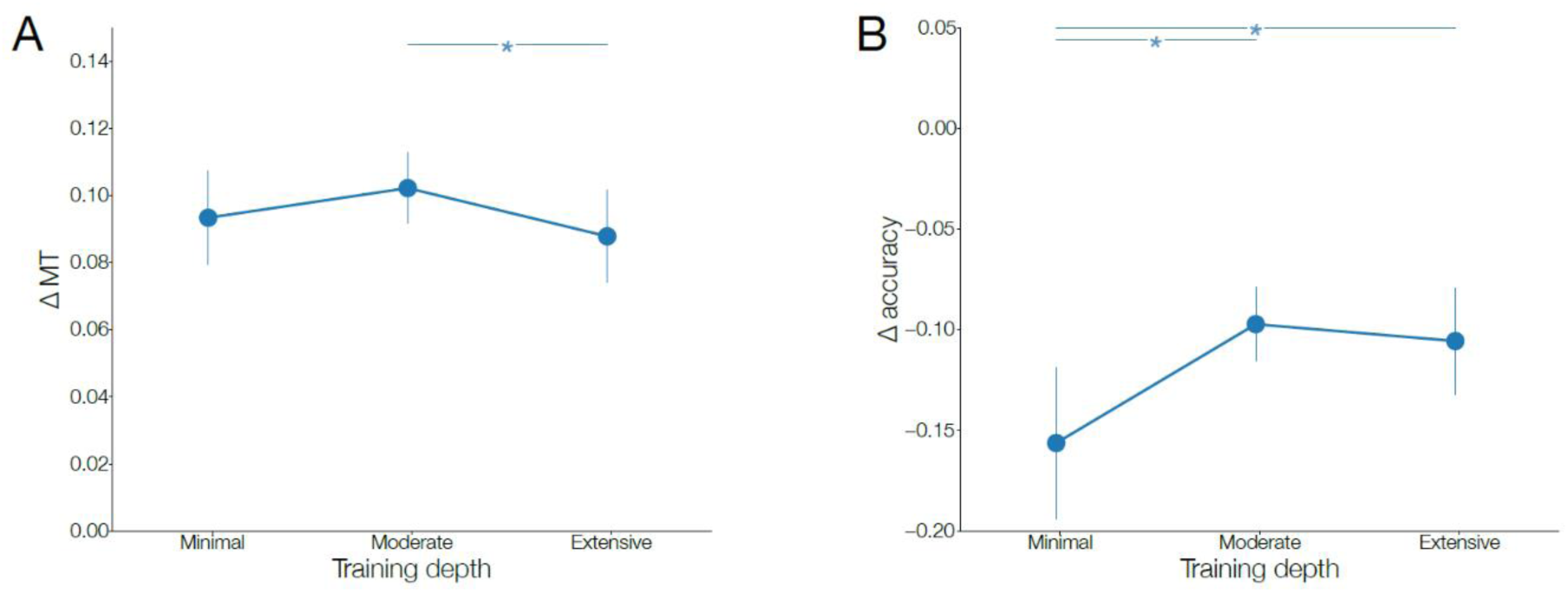
Performance changes following DLPFC stimulation. DLPFC disruption led to deficits in both (A) the speed of execution and (B) the accuracy of execution relative to performance after control stimulation. However, this performance cost was graded by skill level such that the minimally trained skills were impacted the most relative to the moderately and extensively trained skills. ΔMT = % change in movement time relative to control stimulation. Δ accuracy = % change in accuracy relative to control stimulation.

**Figure 6.**
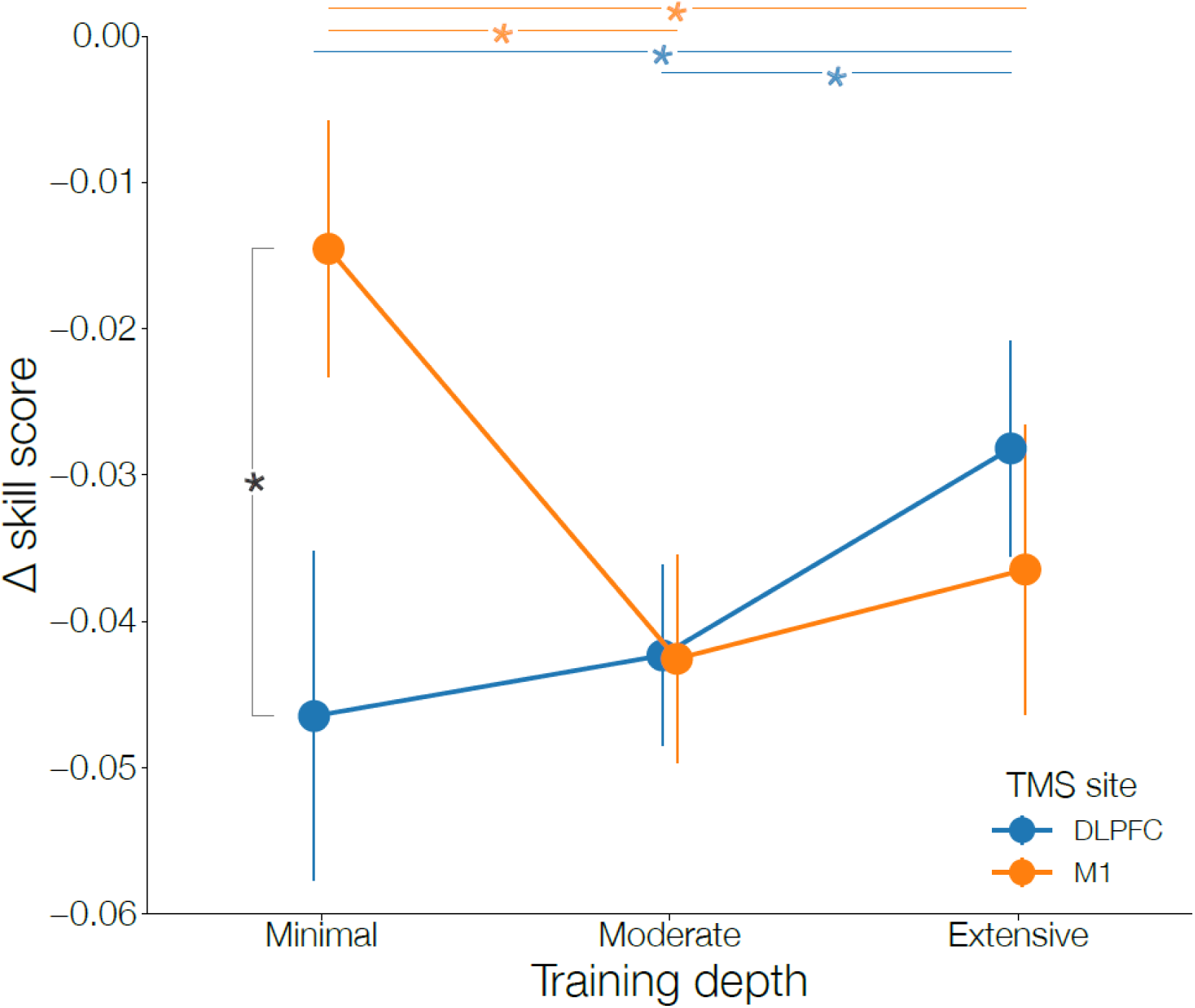
Dissociable impact of M1 and DLPFC stimulation on skilled motor performance as a function of expertise. Although disruption of both M1 and DLPFC led to deficits in skilled performance, we observed a significant interaction between the site of stimulation and the level of training. The cost of disruption was reduced as a function of expertise following DLPFC stimulation but increased as a function of expertise following M1 stimulation.

### Speed-accuracy composite skill score

To control for the well-known tradeoff in speed and accuracy, we calculated composite scores combining both measures to encapsulate the total skill level of our participants for each sequence.

To assess performance for the vertex stimulation condition, we operationalized “skill score” as the number of correct keys pressed per second (KPS). KPS is computed by dividing the total number of key presses after the first key, by the time from the first correct key press to the last key press, regardless of its accuracy. To avoid division by zero, we only included trials with at least 2 key presses in the calculation of KPS.

To examine the impact of DLPFC and M1 stimulation on performance, we took an additional step to incorporate each individual’s baseline level of performance after control stimulation into the skill score. For each participant and each sequence, we computed the average KPS and average partial accuracy in the control session only. Partial accuracy was operationalized as the number of correct key presses after the first key divided by either the number of attempted key presses after the first key, if the trial terminated within 3 s, or 7 otherwise. We then subtracted this baseline from the corresponding KPS or partial accuracy value of each trial of the same sequence in the other TMS sessions of the same participant. This gave us a %Δ*KPS* and %Δ*pacc* value for each trial, both of which scale with the relative change in performance in the same direction (positive = gain, negative = loss). The average of this bivariate measure was taken to index the combined dynamic of speed-accuracy.

## Results

### The evolution of performance during training

Our main hypotheses centered around the roles dorsolateral prefrontal cortex (DLPFC) and primary motor cortex (M1) play in supporting novice, intermediate, and expert motor skills. Participants practiced six separate motor sequences trained to three different depths in 24 separate sessions over a six-week period so that they would all concurrently have two different sequencing skills at each level of expertise.

To confirm that participants successfully learned all sequences during the training period, we compared the raw movement time (MT) required to execute each sequence and the accuracy rate for each sequence at the end of Week 6 to that at the beginning of Week 1. Over the course of training, participants markedly sped up sequence execution as indexed by the total MT required to press all eight keys accurately (Week 6 vs. 1: β = -1183.0, z = -52.8, p < 0.001 overall; p < 0.001 for every skill level). The rate of improvement was higher for more deeply trained sequences compared to the minimally trained ones (*Week* × *Skill level* Extensive vs. Minimal: β = 291.5, z = 4.6, p < 0.001; Moderate vs. Minimal: β = 278.1, z = 4.2, p < 0.001). Participants similarly improved their accuracy rate over the course of training for the moderately and extensively trained sequences (*accuracy* Week 6 vs. 1 Extensive: β = 0.042, z = 3.5, p < 0.001; Moderate: β = 0.068, z = 3.3, p = 0.001). We also observed a quadratic skill level x time interaction suggesting that participants’ accuracy improved to a greater extent on moderately and extensively trained sequences compared to minimally trained sequences (β = 0.052, z = 2.0, p = 0.043).

To evaluate potential ceiling effects toward the end of training, we compared MT at the end of Week 6 to that at the end of Week 5. Highly trained sequences continued to improve even during this last week of training (Week 6 vs. 5 overall: β = -59.5, z = - 2.7, p = 0.008; Extensive: β = -112.0, z = -8.7, p < 0.001; Moderate: β = -127.8, z = -5.6, p < 0.001; Minimal: β = 61.4, z = 1.0, p = 0.323). Note that there were many fewer trials for the minimally trained sequences, which may have limited our power to detect an effect.

We also sought to confirm that participants had three levels of expertise on different sequences concurrently. To this end, we compared MT of the three levels of training at the end of Week 6 in a pairwise manner. In line with our expectations, extensively trained sequences were executed the fastest, followed by moderately trained sequences, and then minimally trained sequences, which were executed the most slowly (Extensive vs. Moderate: β = -324.3, z = -17.5, p < 0.001; Extensive vs. Minimal: β = -1078.6, z = -23.9, p < 0.001; Moderate vs. Minimal: β = -754.4, z = -16.0, p < 0.001). The quadratic contrast across skill level was significant (β = 175.6, z = 7.8, p < 0.001), indicating that the improvement in MT was larger going from minimal to moderate levels of training relative to the improvement observed moderate and extensive training. We observed similar results when examining accuracy rate. Participants were more accurate in completing extensively trained sequences relative to moderately trained sequences (β = 0.041, z = 2.5, p = 0.013) and more accurate on both extensive and moderate sequences when compared with minimally trained sequences (Extensive vs. Minimal: β = 0.314, z = 9.4, p < 0.001; Moderate vs. Minimal: β = 0.273, z = 7.7, p < 0.001). The quadratic contrast across skill level was again significant, indicating that the improvement in accuracy rate was larger going from minimal to moderate levels of training relative to the further improvement seen going from moderate to extensive training (β = -0.095, z = -5.2, p < 0.001).

Individuals learn at different rates, where some improve more rapidly than others during the same training sessions. For example, the extensively trained sequences in some participants may not always be executed within the same range of time as in other participants – although they are always the executed faster than moderate and minimally trained sequences for every participant. To account for such inter-participant variation while maintaining the intraparticipant skill gradient, we standardized the MT separately for each participant by subtracting the participant mean from all values and dividing the differences by the participant standard deviation. This generated a list of z-scored movement times for each participant. Repeating the above analyses on the new metric yielded the same results as observed in the raw MT when comparing improvement over the course of training and the levels of skill attained at the end of training (Week 6 vs. 1: β = -1.714, z = -54.0, p < 0.001 overall; p < 0.001 for every skill level; *Week* × *Skill level* Extensive vs. Minimal: β = 0.287, z = 3.2, p = 0.001; Moderate vs. Minimal: β = 0.277, z = 3.0, p = 0.003; Week 6 vs. 5: β = -0.085, z = -2.7, p = 0.007 overall; p < 0.001 for Extensive and Moderate levels, p = 0.194 for Minimal level; end-of-training Extensive vs. Moderate: β = -0.475, z = -18.2, p < 0.001; Extensive vs. Minimal: β = -1.586, z = -24.8, p < 0.001; Moderate vs. Minimal: β = -1.111, z = -16.7, p < 0.001; Linear contrast across skill levels: β = -1.122, z = -24.8, p < 0.001; Quadratic contrast: β = 0.260, z = 8.1, p < 0.001) (**Figure 3a**).

### Effects of training (i.e., performance following control stimulation)

Our active control stimulation condition serves as a comparison for the effects of DLPFC and M1 stimulation as it controls for nonspecific effects of stimulation (e.g., auditory clicks, sensation of stimulation at the scalp, current induced in the cortex, etc.). As such, we first verified that the basic pattern of results from the last training session still held for all participants in this session. We confirmed the effect of training depth on standardized MT, where the more deeply trained sequences were executed more quickly than the more shallowly trained sequences during this session (Linear contrast across skill levels: β = -0.868, z = -35.2, p < 0.001; Quadratic contrast: β = 0.108, z = 4.5, p < 0.001; Extensive vs. Moderate: β = -0.5, z = -14.5, p < 0.001; Extensive vs. Minimal: β = -1.2, z = -35.2, p < 0.001; Moderate vs. Minimal: β = -0.7, z = -21.4, p < 0.001). We also observed a significant effect of training depth on accuracy, where the more deeply trained sequences were executed more accurately than the minimally trained sequences (Linear contrast across skill levels: β = 0.494, z = 3.2, p < 0.001; Quadratic contrast: β = -0.354, z = -2.3, p = 0.022; Extensive vs. Minimal: β = 0.698, z = 3.2, p < 0.001; Moderate vs. Minimal: β = 0.783, z = 3.6, p < 0.001). We did not observe a significant difference in accuracy between extensively trained sequences and moderately trained sequences (Extensive vs. Moderate: β = -0.084, z = -0.4, p = 0.698). Accuracy on extensively and moderately trained sequences potentially converged due to ceiling effects (Extensive accuracy rate = 84.7%, moderate accuracy rate = 85.9%, minimal accuracy rate = 72.1%). These results indicate that extensive training did in fact improve performance and does not reflect a simple shift along a single speed-accuracy function as sequences could be performed more quickly without an increase in errors (**Figure 3b,c**).

Given the coupled improvement in both movement time and accuracy as a function of training depth, we sought to unify these quantities into a single metric to encapsulate performance at each level of skill (see Methods). For this purpose, we calculated a “skill score” that combines the number of correct keys pressed as a function of time to assess the combined effect of movement time and accuracy: Faster movement time coupled with higher accuracy should strongly increase skill score, and vice versa. We again found a main effect of training depth on skill score where more keys were pressed correctly within a shorter amount of time for deeply trained sequences compared to shallowly trained ones (Linear contrast across skill levels: β = 0.786, z = 27.6, p < 0.001; Quadratic contrast: β = -0.059, z = -2.1, p = 0.040; Extensive vs. Moderate: β = 0.48, z = 12.0, p < 0.001; Extensive vs. Minimal: β = 1.11, z = 27.6, p < 0.001; Moderate vs. Minimal: β = 0.63, z = 15.5, p < 0.001).

One possible mechanism by which training can improve performance is via an improvement in motor planning (rather than in simply execution itself). Recall that each sequence is cued in advance and there is a variable delay between cue presentation and the presentation of the first item in the sequence. Thus, one index of motor planning is the initial response time (initiation time) to the presentation of the first item in each sequence as this first key press could be prepared and deployed immediately as the associated key position was illuminated. This hypothesis found some support as initiation time was significantly faster for the extensively trained sequences compared to the moderately and minimally trained sequences (Extensive vs. Moderate: β = -21.1, z = -4.5, p < 0.001; Extensive vs. Minimal: β = -22.3, z = -4.6, p < 0.001). Though numerically in the same direction, we did not observe a significant effect when comparing moderately to minimally trained sequences (β = -1.2, z = -0.3, p = 0.801). Furthermore, key-to-key transitions also benefited from training as we observed a main effect of training depth for all key positions (p < 0.001 for all pairwise comparisons at all transitions, except for Extensive vs. Moderate at the 2-to-3 transition, p = 0.013). These results suggested that training had beneficial impacts in all aspects of the motor skill, from planning through execution.

### Effects of M1 disruption

BOLD activity in M1 has been reported to decrease over the course of training during the performance of motor sequencing skills(Dayan & Cohen, 2011; Steele & Penhune, 2010). Some previous researchers have made the conjecture that this decrease reflects an increase in metabolic efficiency of the region as learning progresses (Kornysheva & Diedrichsen, 2015; Steele & Penhune, 2010; Wymbs & Grafton, 2015). However, other recent neuroimaging studies have struggled to find any evidence for distinct skill-specific patterns of activity in M1 as skill develops(Yokoi et al., 2018; Yokoi & Diedrichsen, 2019). The evolving causal role for M1 in supporting motor skill as a function of expertise has thus remained unclear. Our design allows us to make causal inferences regarding the role of M1 in motor skills as we disrupt M1 activity with inhibitory TMS and examine its impact on behavioral performance at each level of skill.

M1 disruption produced deficits in the speed of execution for all skill levels relative to the control stimulation condition (%Δ*MT* Minimal: β = 0.047, z = 2.2, p = 0.029; Moderate: β = 0.097, z = 4.6, p < 0.001; Extensive β = 0.092, z = 4.3, p < 0.001). We next examined whether the disruptive effects of stimulation varied as a function of skill level. We found strong evidence for an effect of skill level on the impact of M1 disruption. M1 stimulation produced larger impairments in performance for the more extensively trained sequences (Linear contrast across skill levels: β = 0.032, z = 7.3, p < 0.001; Quadratic contrast: β = -0.023, z = -5.4, p < 0.001) (**Figure 4a**). Extensively and moderately trained sequences were executed more slowly than minimally trained sequences after controlling for their baseline differences ((%Δ*MT M*1 *vs*. *vertex* Extensive vs. Minimal: β = 0.045, z = 7.3, p < 0.001; Moderate vs. Minimal: β = 0.050, z = 8.2, p < 0.001). This effect of M1 TMS on movement time cannot be attributed to a speed-accuracy tradeoff. Accuracy rate was either unaffected (%Δ*acc M*1 *vs*. *vertex* Minimal: β = -0.050, z = -1.6, p = 0.104) or significantly worse following M1 stimulation (Extensive: β = -0.090, z = -2.9, p =0.004; Moderate: β = - 0.115, z = 3.7, p < 0.001). We found some evidence that accuracy rate was more impaired for the higher levels of training, confirming that the deleterious impact of M1 stimulation is greater at higher levels of skill (%Δ*acc M*1 *vs*. *vertex* Linear contrast across skill level: β = -0.028, z = -1.8, p = 0.074; Quadratic contrast: β = 0.037, z = 2.4, p = 0.018; Extensive vs. Moderate: β = 0.025, z = 1.2, p = 0.250; Extensive vs. Minimal: β = -0.040, z = -1.8, p = 0.074; Moderate vs. Minimal: β = -0.065, z = -2.9, p = 0.003) (**Figure 4b**).

Our combined speed-accuracy skill score similarly reflects that the causal importance of M1 in supporting skilled performance grows as a function of expertise. M1 stimulation negatively impacted both extensively trained and moderately trained skills to a greater extent than minimally trained skills (Δ*skill score M*1 *vs*. *vertex* Linear contrast across skill level: β = -0.016, z = -4.2, p < 0.001; Extensive vs. Minimal: β = -0.022, z = - 4.2, p < 0.001; Moderate vs. Minimal: β = -0.028, z = -5.4, p < 0.001; Extensive vs. Moderate: β = 0.006, z =1.2, p = 0.243) (**Figure 6, orange curve**).

### Effects of DLPFC disruption

Similar to M1, BOLD activity in DLPFC has been reported to decrease over the course of motor skill training. Unlike M1, however, this finding has usually been interpreted as a reduced role for the region in supporting expert motor skills. If true, we would expect that DLPFC stimulation would have a large impact on performance for minimally trained skills and a negligible impact on the most extensively trained skills.

We found that disruption of DLPFC produced behavioral performance deficits for all levels of expertise. The speed of execution was impaired relative to the control stimulation condition for all skill levels (%Δ*MT DLPFC vs*. *vertex* Minimal: β = 0.093, z = 4.4, p < 0.001; Moderate: β = 0.103, z = 4.9, p < 0.001; Extensive: β = 0.091, z = 4.3, p < 0.001). Accuracy also suffered at all skill levels when this site was stimulated (%Δ*acc DLPFC vs*. *vertex* Minimal: β = -0.157, z = -5.0, p < 0.001; Moderate: β = -0.096, z = -3.1, p = 0.002; Extensive: β = 0.106, z = -3.4, p < 0.001).

We next queried whether this disruptive effect of DLPFC stimulation varied as a function of skill level. Stimulation had a larger negative impact on normalized MT for moderately trained sequences relative to the extensively trained sequences (%Δ*MT DLPFC vs*. *vertex* Extensive vs. Moderate: β = -0.012, z = -2.0, p = 0.046) (**Figure 5a**). We did not observe significant differences in the impact of stimulation when comparing minimally trained sequences to either of the other two levels of skill (%Δ*MT DLPFC vs*. *vertex* Extensive vs. Minimal: β = -0.002, z = -0.3, p = 0.744; Moderate vs. Minimal: β = 0.010, z = 1.5, p = 0.123). However, when examining the disruptive effect of DLPFC stimulation on accuracy rate, we observed a significant linear trend across skill level (%Δ*acc DLPFC vs*. *vertex* β = 0.036, z = 2.2, p = 0.028) (**Figure 5b**). The impact of DLPFC stimulation was larger for minimally trained sequences than in the other, more highly trained sequences (Extensive vs. Minimal: β = 0.050, z = 2.2, p = 0.028; Moderate vs. Minimal: β = 0.060, z = 2.6, p = 0.008). We did not observe a difference in the impact of stimulation on accuracy between extensively trained and moderately trained sequences (Extensive vs. Moderate: β = -0.010, z = -0.04, p = 0.664).

The effects of DLPFC stimulation on execution speed and accuracy rate produced a mixed pattern of results across skill level. However, this is the precise situation in which our composite speed-accuracy skill score can be illuminating. We observed a significant linear trend across skill level (β = 0.014, z = 3.6, p < 0.001), suggesting that the impact of DLPFC stimulation diminished as a function of expertise. Specifically, the deleterious effect of DLPFC stimulation on skill score was significantly larger for minimally trained skills when compared to the more extensively trained skills (Extensive vs. Minimal: β = 0.019, z = 3.6, p < 0.001). The same was true for moderately trained skills relative to extensively trained skills (Extensive vs. Moderate: β = 0.014, z = 2.6, p = 0.008) (**Figure 6, blue curve**). We did not find a significant difference in the impact of stimulation on skill score between moderately trained and minimally trained sequences (Moderate vs. Minimal: β = 0.005, z = 0.9, p = 0.352). This pattern of results all point to the fact that DLPFC disruption had a graded impact on performance whereby the effect of stimulation was more muted for the most extensively trained skills. However, we found that DLPFC stimulation led to deficits in performance as measured by this composite skill score for all skills regardless of the level of training (Minimal: β = -0.048, z = -3.8, p < 0.001; Moderate: β = -0.043, z = -3.4, p = 0.001; Extensive: β = -0.028, z = -2.3, p = 0.022).

### The effects of DLPFC and M1 disruption on performance are dissociable

The above results establish that both DLPFC and M1 stimulation had a deleterious impact on skilled motor performance and that disruption of both sites showed a graded impact as a function of skill level. Are the effects of stimulation between the two sites dissociable as a function of skill level? We found strong evidence for just such a dissociation.

TMS site significantly modulated the effect of skill level on normalized movement time (MT) (site x skill level interaction for %Δ*MT* Linear contrast: β = -0.033, z = -5.3, p < 0.001). We observed a similar significant interaction when examining the impact of stimulation on accuracy rate (site x skill level interaction for %Δ*acc* Linear contrast: β = 0.064, z = 2.8, p = 0.005; Quadratic contrast: β = -0.066, z = -2.9, p = 0.004), suggesting that the dissociable effect of stimulation to DLPFC and M1 could not simply be attributed to a speed-accuracy tradeoff.

We then directly compared the effect of stimulation on our composite skill score across the two stimulation sites of interest. Overall, DLPFC stimulation had a larger negative impact on skill score than M1 stimulation did (Δ*skill score* β = -0.008, z = -2.7, p = 0.007). However, we found a significant interaction between the effect of TMS site and training depth on the composite skill score (site x skill level interaction for Δ*skill score* β = 0.029, z = 5.5, p < 0.001) (**Figure 6**). This interaction was driven by the fact that there was a much larger harmful impact of DLPFC stimulation on minimally trained skills relative to M1 stimulation (Minimal: β = -0.033, z = -6.2, p < 0.001; Moderate: β = -0.000, z = -0.02, p = 0.985; Extensive: β = 0.008, z = 1.5, p = 0.132). The causal effect of DLPFC and M1 disruption on motor performance depends on the proficiency that an individual has with the executed sequence.

## Discussion

We trained human participants on six separate motor sequences at three levels of training intensity (minimal, moderate, extensive) over a six-week training period. Although BOLD activity in both dorsolateral prefrontal cortex (DLPFC) and motor cortex (M1) has been reported to decrease as expertise develops, this has been assumed to reflect a reduced role for DLPFC in supporting performance but increased efficiency for M1. Here, we causally test this assumption by applying inhibitory rTMS over each site just prior to an assessment of performance at each skill level. Although cortical disruption at both sites produced deficits in performance for all motor skills relative to a control stimulation condition, we found dissociable effects on skilled performance as a function of skill level. Specifically, DLPFC stimulation led to the greatest cost for minimally trained skills and relatively smaller costs to performance for the moderately and extensively trained skills. M1 stimulation led to the opposite pattern of results whereby performance on the more extensively trained skills was impacted the most with relatively small costs for the minimally trained skills. These results confirm the assumptions of many models of motor skill acquisition, but also point to a maintained role for DLPFC in supporting performance even for the most highly trained skills.

Over the six-week training period, participants improved both the speed and accuracy of performance across all motor sequences as expected. We validated our within-participant experimental design where, at the end of the training, each individual demonstrated a performance gradient across the extensively-, moderately-, and minimally-trained sequences. This design had the advantage of allowing us to make inferences about the neural mechanisms supporting each skill level within a single session without temporal confounds (e.g., general task familiarity) that might arise when using a single sequence examined at different time points during skill development. Although several studies have queried the role of both DLPFC and M1 in motor sequencing skills with non-invasive brain stimulation (CITE), to our knowledge, the present study is the first to examine the impact of stimulation following extensive training on a novel motor skill. Most prior work has either examined the effects of cortical disruption on skill acquisition in a single session (Pascual-Leone et al., 1996; Robertson et al., 2001; Wilkinson et al., 2010) or has interrogated how stimulation early on in training affects consolidation and/or retrieval after long delays (Cohen & Robertson, 2011; Galea et al., 2010; Greeley et al., 2020; Reis et al., 2009). Our results provide the first causal evidence in humans for the evolving roles of both M1 and DLPFC over the course of long-term skill development.

Our finding that DLPFC plays a causal role in motor sequencing performance on novice skills mirrors the results from several studies, however, most of these studies were concerned with initial skill acquisition rather than the expression of a well-learned skill. Several groups have found that inhibitory TMS over DLPFC impairs performance and disrupts sequence learning in a single session (Dayan et al., 2018; Robertson et al., 2001, 2001). It is important to note that Robertson and colleagues (2001) found this was only true for spatially cued sequences but not when movements needed to be learned based on symbolic cues presented in a single location. Patients with lesions to prefrontal cortex also display sequence learning deficits on motor sequencing tasks (Beldarrain et al., 2002). More recently, Greeley et al. (2020) found that applying transcranial direct current stimulation (tDCS) over DLPFC similarly has detrimental impacts on sequence learning regardless of the polarity of stimulation. Participants in our study had initial exposure to the minimally trained sequences and showed improvement in performance over the course of training prior to the first stimulation session. Similar to the studies cited above, we likewise found that disruption of DLPFC led to impaired performance for these novice skills. Several studies have suggested that DLPFC activity may actually be detrimental to the development and execution of trained motor skills (Ambrus et al., 2020; Cohen & Robertson, 2011; Galea et al., 2010). Non-invasive brain stimulation that disrupts DLPFC activity following a bout of training can lead to benefits in consolidation and improved performance on subsequent sessions (Cohen & Robertson, 2011; Galea et al., 2010). There is also neuroimaging evidence that individuals who improve the most during long-term motor sequence training are those who display a release of prefrontal cortex functional connectivity with sensorimotor areas (Bassett et al., 2015). Our results, however, suggest that DLPFC activity is causally necessary for successful performance, even for extensively trained skills. Given the purported role of DLPFC in supporting working memory operations in motor skills, one might expect the cost to performance for the extensively trained skills to be related to impairments in the maintenance of the sequence cue over the brief delay just before executing the sequence. However, we found no difference in the time it takes to initiate sequences between the stimulation conditions, suggesting that the cost to performance following DLPFC stimulation comes during skill execution.

The role of M1 in supporting skilled motor performance has come into question in recent years. Non-invasive brain stimulation studies suggest that disrupting M1 activity impairs initial skill acquisition and novice performance, but our findings indicate a relatively minimal impact on novice skills. However, recent work with rodents has suggested that while M1 may be necessary for the acquisition of a simple motor skill, it plays a minimal role in the execution of a skill once it has been trained extensively (Kawai et al., 2015). Along a similar vein, several neuroimaging studies have reported that M1 may not be involved in supporting motor sequencing skills beyond the simple execution of each individual motor element in a sequence (Berlot et al., 2020; Kornysheva & Diedrichsen, 2015; Yokoi et al., 2018). These studies employ representational similarity analyses (RSA) and other multivariate analyses to attempt to identify brain areas that contain representations of trained motor sequences. These representations have been very difficult to read out from primary motor cortex and seem to reside in more anterior premotor areas in frontal cortex (e.g., SMA, pre-SMA, premotor cortex). This finding has been puzzling and has left it unclear whether M1 is only important for individual low-level motor commands, whether there is a limit to the inferences that can be drawn from multivariate fMRI analyses, or perhaps whether the skill that is learned in motor sequencing tasks is more of a ‘cognitive’ rather than a motor skill (Wong & Krakauer, 2019). If M1 does not contribute to the skill itself and only in the execution of each individual element, we would have expected that rTMS over M1 would lead to a relatively static cost to performance for all levels of skill. However, given that the cost of cortical disruption grows larger as expertise develops, our results indicate that M1 plays an integral role in supporting expert motor sequencing skills in line with prior work (Floyer-Lea, 2004; Karni et al., 1998; Penhune & Doyon, 2002).

Several models of skill development have suggested that the initial learning of a skill relies on more cognitive “higher-level” processing that becomes less necessary over time (Doyon & Benali, 2005; Fitts & Posner, 1967; Hikosaka et al., 2002; Verwey et al., 2015; Willingham, 1998). For example, the Cognitive framework for Sequential Motor Behavior (C-SMB) that was developed with a particular eye toward the DSP task used here posits that motor sequencing skills require coordination between a central processor presumed to be supported by DLPFC and a motor processor involving M1 (Verwey et al., 2015, 2019). Motor skills are initially based on a central-symbolic mode that relies on cognitively demanding spatial and verbal representations supported by the central processor. After extensive practice, motor elements are chunked together into motor sequence representations that can be executed by a motor processor that needs minimal input from the central processor following action selection. Similarly, Hikosaka and colleagues proposed that early motor sequence learning requires transformation between spatial and motor coordinates that requires a frontoparietal associative network (2002). However, later stages of learning and skill expression are dominated by slow-learning sensorimotor circuits, as performance can rely on a motor sequencing mechanism without the need for a spatial-motor transformation. Our results are largely in line with these models as the detrimental impact of DLPFC stimulation diminishes as expertise develops while the detrimental impact of M1 stimulation becomes more pronounced. The fact that DLPFC stimulation still carries performance costs even for the most highly trained skills is perhaps most consistent with the C-SMB, which suggests that chunked motor representations must still be selected and loaded by a central processor prior to execution. However, an alternate view would be that DLPFC stimulation affects expert performance only because the DSP task we employed has a high demand on visuospatial processing. As noted above, there is some evidence to suggest that DLPFC activity is more involved in motor sequencing tasks when movements are spatially cued relative to symbolically cued movements (Bo et al., 2011; Koch & Hoffmann, 2000; Robertson et al., 2001). Future work is necessary to disentangle whether DLPFC’s maintained role in supporting expert motor skills is specific to tasks with a visuospatial component.

In summary, the present study provides causal evidence of the dissociable roles of DLPFC and M1 over the course of long-term motor skill development but shows that both regions are instrumental in supporting highly trained skills. Our findings support models of skill learning that posit separable associative and sensorimotor systems that display different time courses of involvement over the course of learning, but also suggest that both systems are required for performance at all levels of skill.

## Acknowledgments

We would like to thank James A. Brissenden, Joseph Deluisi, and Michael Vesia for their helpful comments during the drafting of this manuscript

